# DeepEMhancer: a deep learning solution for cryo-EM volume post-processing

**DOI:** 10.1101/2020.06.12.148296

**Authors:** R Sanchez-Garcia, J Gomez-Blanco, A Cuervo, JM Carazo, COS Sorzano, J Vargas

## Abstract

Cryo-EM maps are valuable sources of information for protein structure modeling. However, due to the loss of contrast at high frequencies, they generally need to be post-processed to improve their interpretability. Most popular approaches, based on B-factor correction, suffer from limitations. For instance, they ignore the heterogeneity in the map local quality that reconstructions tend to exhibit. Aiming to overcome these problems, we present DeepEMhancer, a deep learning approach designed to perform automatic post-processing of cryo-EM maps. Trained on a dataset of pairs of experimental maps and maps sharpened using their respective atomic models, DeepEMhancer has learned how to post-process experimental maps performing masking-like and sharpening-like operations in a single step. DeepEMhancer was evaluated on a testing set of 20 different experimental maps, showing its ability to obtain much cleaner and more detailed versions of the experimental maps. Additionally, we illustrated the benefits of DeepEMhancer on the structure of the SARS-CoV-2 RNA polymerase.

## Introduction

Almost one decade after the beginning of the so-called “resolution revolution”, cryogenic electron microscopy (cryo-EM) has become one of the most versatile tools in the field of structural biology. Beginning from thousands of single particle projection images, cryo-EM workflows are capable of obtaining three-dimensional (3D) reconstructions of many macromolecules at “near-atomic” resolution levels. However, the ultimate goal of cryo-EM Single Particle Analysis is not the obtention of 3D maps but the detailed atomic understanding through the derivation of atomic models.

During the atomic model building process, raw 3D maps are rarely employed, as they suffer from loss of contrast at high resolution (Rosenthal and Henderson, 2003) that makes difficult the detection and interpretability of residues and secondary structure. Fortunately, loss of contrast can be alleviated using different contrast restoration algorithms, which are usually known as sharpening methods. The first sharpening approach for cryo-EM maps was introduced by Rosenthal and Henderson (Rosenthal and Henderson, 2003) and its formulation, based on the B-factor correction, is still at the basis of the most commonly employed sharpening methods, including RELION postprocessing (Kimanius et al., 2016; Zivanov et al., 2018) or Phenix AutoSharpen (Terwilliger et al., 2018). The principle behind these algorithms consists in the correction of the raw maps by boosting the amplitude of their high frequency Fourier components. The strength of the amplitude boost at each frequency depends on the frequency itself and on a single number, the B-factor, that measures the global loss of contrast. Thus, although the different B-factor-based methods differ in the procedures employed to determine the B-factor that is applied, they modify the volume globally in a similar manner.

Despite being widely used, B-factor-based approaches present an important limitation: they do not consider the differences in quality that different parts of the map may present and they produce density maps that do not correspond to the scattering properties of biological macromolecules (Vilas et al., 2020). Consequently, for the case of maps that exhibit heterogeneous local resolution, some regions could be undersharpened whereas others could be oversharpened. Recently, local sharpening algorithms, that alleviate this shortcoming, have been proposed. Thus, the LocScale (Jakobi et al., 2017) algorithm uses the information contained in an atomic model to locally scale up a map. Such transformation is achieved by means of a sliding window approach in which the amplitudes of the map region that lay inside the window are scaled up to agree with the atomic model provided. Following a totally different strategy, the LocalDeblur (Ramírez-Aportela et al., 2020) algorithm employs a Wiener filtering approach that performs local deblurring with a strength proportional to an estimation of the local resolution, that has to be pre-computed. Similarly, LocSpiral (Kaur et al., 2020) employs the spiral phase transformation to factorize the volume and then, perform a local enhancement based on the normalization and thresholding of the amplitudes.

Despite their benefits, current local sharpening approaches present some drawbacks. Thus, both LocSpiral and LocalDeblur depend on masks to distinguish the macromolecule from the noise and LocalDeblur requires also from an estimation of the local resolution of the map. On the other hand, the main strength of LocScale, its ability to employ the structural information of atomic models, could also be regarded as its main weakness since the availability of atomic models limits its applicability.

With the aim of overcoming these shortcomings, in this work, we present Deep cryo-EM Map Enhancer (DeepEMhancer), a fully automatic deep learning-based approach that performs cryo-EM volume post-processing. Deep learning has revolutionized the field of Artificial Intelligence and its impact has been felt in many others including cryo-EM. Deep learning in cryo-EM was firstly applied for the problem of particle picking (Wagner et al., 2019; Wang et al., 2016; Zhu et al., 2017) and since then, it has evolved to deal with other questions such as map reconstruction (Gupta et al., 2020; Zhong et al., 2019), map segmentation (Maddhuri Venkata Subramaniya et al., 2019; Si et al., 2020) or local resolution determination (Avramov et al., 2019; Ramírez-Aportela et al., 2019). As in most of those methods, our approach relies on a convolutional neural network (CNN) that is trained on massive quantities of data. Particularly, our development, that follows a simple image super-resolution setup (Yang et al., 2019), exploits the vast amount of structural information that is contained in the Electron Microscopy Data Bank (EMDB) database (Lawson et al., 2015) in order to mimic the local sharpening effect of the LocScale algorithm. However, DeepEMhancer does not require any atomic model to function and, contrary to previous methods, it also performs automatic (tight) masking of input maps. Our results show that DeepEMhancer, that works in a fully automatic manner, is able to largely improve the interpretability of the maps contained in our benchmark, performing better than classical B-factor approaches.

## Results

DeepEMhancer is based on an end-to-end U-net architecture (Ronneberger et al., 2015) trained in a supervised manner. Particularly, we implemented a 3D U-net consisting in three downsampling blocks and three upsampling blocks that processes cubic chunks of the input map. Training was performed using pairs of input maps and target maps, consisting in experimental cryo-EM maps and tightly masked LocScale post-processed maps. See Methods section for a complete description of the data preparation, training, and evaluation processes.

### Deep cryo-EM Map Enhancer performance on the testing set

In order to assess the quality of DeepEMhancer predictions, we first compared them against the target maps generated by LocScale. Thus, for DeepEMhancer maps, we measured a median correlation coefficient of 0.91 against LocScale maps in contrast to 0.60 for input maps (see Supplementary Material Figure S1). Such an important increase in the correlation coefficient implies that DeepEMhancer has learned to accurately reproduce the effect of LocScale sharpening with one important advantage: no atomic models are required to employ DeepEMhancer.

Although reproducing the LocScale sharpening effect was our main objective, the ultimate goal of map post-processing is to simplify the process of atomic model building. With the aim of studying if DeepEMhancer also contributes to that purpose, we next explored whether DeepEMhancer post-processed maps were more similar to the actual atomic models. To do so, we computed, for all the maps included in the testing set, the Fourier Shell

Correlation coefficient (FSC) resolution between the input (half maps average) and post-processed maps against the reference maps obtained from the atomic models. As it is shown in Figure 1, for all the examples included in the testing set, the application of DeepEMhancer increased the similarity of the input maps with respect to the references (blue and green bars). Indeed, the post-processed maps exhibit a median FSC resolution value of 3.3 Å compared to 3.9 Å for the input maps. Particularly, the median improvement achieved by DeepEMhancer was ∼0.6 Å (∼14% in the frequency domain). Such an important improvement confirms that the maps computed by DeepEMhancer are much more similar to the target maps.

**Figure 1.**
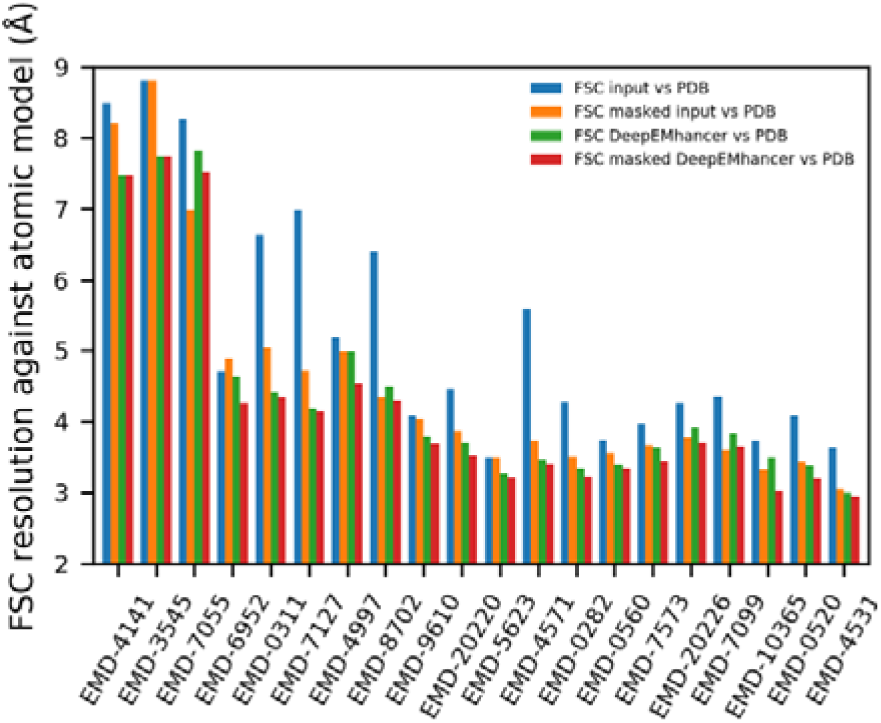
DeepEMhancer produces maps that are more similar to the atomic models. Resolution (determined by Fourier Shell Correlation coefficient, FSC) between the reference maps obtained from the atomic model and 1) the input maps (blue), 2) the input maps tightly masked (orange), 3) the post-processed maps by DeepEMhancer (green) and 4) the post-processed maps by DeepEMhancer tightly masked (red). EMDB entries are sorted by published global resolution.

DeepEMhancer post-processing operation performs a non-linear transformation of the experimental volume that produces a set of effects that could be broadly classified as masking/denoising and sharpening-like features enhancement. In order to disentangle the contribution of the different effects, we have also computed the FSC of the input and post-processed maps using a very tight mask derived from the atomic model. As it can be observed in Figure 1, the FSC resolution obtained for the post-processed maps tend to be better than the values computed for the input independently of the mask application (green and red bars vs orange bar), which implies that the masking effect is of high-quality, as the resolutions for the unmasked DeepEMhancer results tend to be better than the ones for the masked input maps.

Similarly, and, although it is true that the trend is not as strong as in the previous experiment, DeepEMhancer also tends to improve the resolution of the masked regions (Figure 1, orange vs red bars), which supposes an enhancement of the map features. Leaving aside some problematic examples such as EMD-7055 (Tenthorey et al., 2017), that will be discussed in Supplementary Material, most of the evaluated maps exhibit a non-negligible improvement in resolution, especially notable when compared to B-factor-based results (see next section), with a median value of ∼0.3 Å.

Alternatively, with the aim of obtaining a complementary measurement of improvement, we computed the DeepRes local resolution for the input and post-processed maps. As can be appreciated in Figure 2, all test cases treated with DeepEMhancer improved in terms of DeepRes local resolution, with dramatic improvements of more than 0.8 Å and a median improvement of ∼0.4 Å. Again, those figures, consistent with the FSC-based measurements, point out that DeepEMhancer is improving the interpretability of the maps.

**Figure 2.**
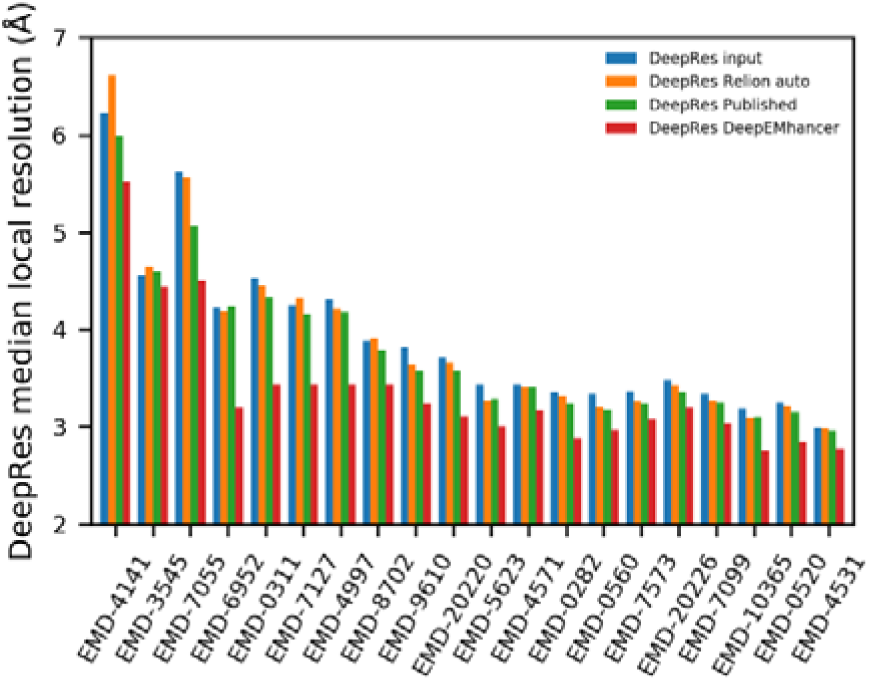
DeepEMhancer produces better quality maps. DeepRes median local resolution estimation for 1) the input maps (blue), 2) the post-processed maps obtained with Relion postprocessing automatic B-factor (orange), 3) the post-processed maps deposited in EMDB (green) and 4) the post-processed maps obtained with DeepEMhancer (Red). EMDB entries are sorted by published global resolution.

### Comparison with B-factor-based methods

With the aim of comparing DeepEMhancer with the commonly employed B-factor-based sharpening methods, we repeated the same experiments using the post-processed maps obtained with the Relion postprocessing algorithm (Kimanius et al., 2016; Zivanov et al., 2018). Before, it is important to notice that contrary to DeepEMhancer, Relion automatic masking is a simple process and thus, in order to make the comparison more interesting, we used instead the masks derived from the atomic models.

Still, when we evaluated the FSC for the masked regions, only a few maps improved, while many others worsened, leading to a median improvement that was negligible (<0.05 Å) for both FSC and median DeepRes resolution (see Figure 2 and Figure 3).

**Figure 3.**
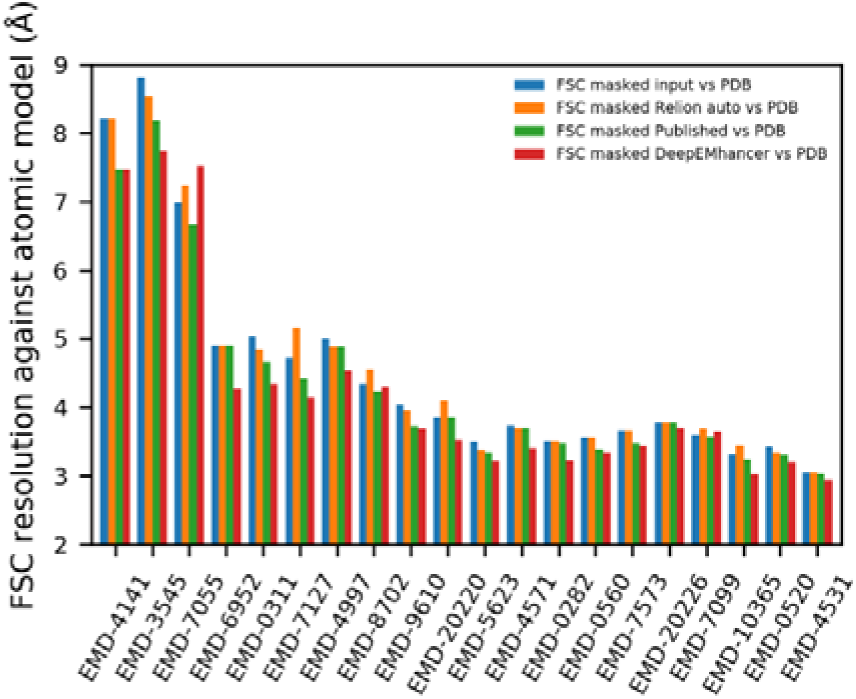
DeepEMhancer produces better results than B-factor-based methods. Resolution (determined by Fourier Shell Correlation coefficient, FSC) between the reference maps obtained from the atomic model and 1) the input maps (blue), 2) the post-processed maps obtained with Relion postprocessing automatic B-factor (orange), 3) the post-processed maps deposited in EMDB (green) and 4) the post-processed maps obtained with DeepEMhancer (red). EMDB entries are sorted by published global resolution.

We acknowledge that the automatic determination of the B-factor can lead to less accurate results than if it were manually selected and it may be the reason behind the poor observed performance. Thus, we have also included in the comparison the post-processed maps deposited in EMDB in which the estimation of B-factor was carried out by the authors. In this case, the improvement in resolution, with median values of ∼0.15 Å and ∼0.1 Å for DeepRes and FSC respectively, although closer to the values obtained using DeepEMhancer, are still considerably inferior (see Figure 2 and 3). In the light of these results, we can state that DeepEMhancer maps tend to be much more similar to the atomic models than the ones obtained using B-factor-based methods and thus, more useful for the process of model building.

### Visual inspection of testing maps

The purpose of this section is to further explore the results obtained with DeepEMhancer for some of the maps included in the testing set with the aim of illustrating how the improvements in global quality measurements translate to tangible improvements in the quality of the maps.

#### EMD-7099

The EMD-7099 (Johnson and Chen, 2018) is a high-resolution volume (global resolution 3.1 Å) of a multidrug resistance ATP-driven pump. EMD-7099 presents 17 transmembrane helices and, although the overall quality of the map is excellent, visualizing the transmembrane regions is challenging because of the signal that comes from the lipids. As a result, important parts of the protein are not traced. Due to the fact that DeepEMhancer was trained to ignore the signal coming from lipidic layers, this example illustrates the unique characteristics of DeepEMhancer when applied to membrane proteins. Thus, as can be observed in Figure 4a-d, DeepEMhancer has been able to suppress the signal coming from the lipid layer in a much more simple and effective way than diminishing the threshold in the raw map or the B-factor-based sharpened maps. The noise suppression effect simplifies the process of model building, as the researchers do not have to deal with masks or larger thresholds that make the visualization of near to noise level features more difficult. Yet not only DeepEMhancer produces a noise reduction effect, but also it is able to enhance some parts of the map that under B-factor based sharpening seem noisy and disconnected. Such improvement, although observed in several regions of the map, is more noticeable at the transmembrane region Thus, the most important enhancement is depicted in Figure 4e-f, in which an important part of the backbone of the protein has been *de novo* traced thanks to DeepEMhancer enhancement, that has restored the densities corresponding to residues A195 to I203 in chain A of PDB 6bhu.

**Figure 4.**
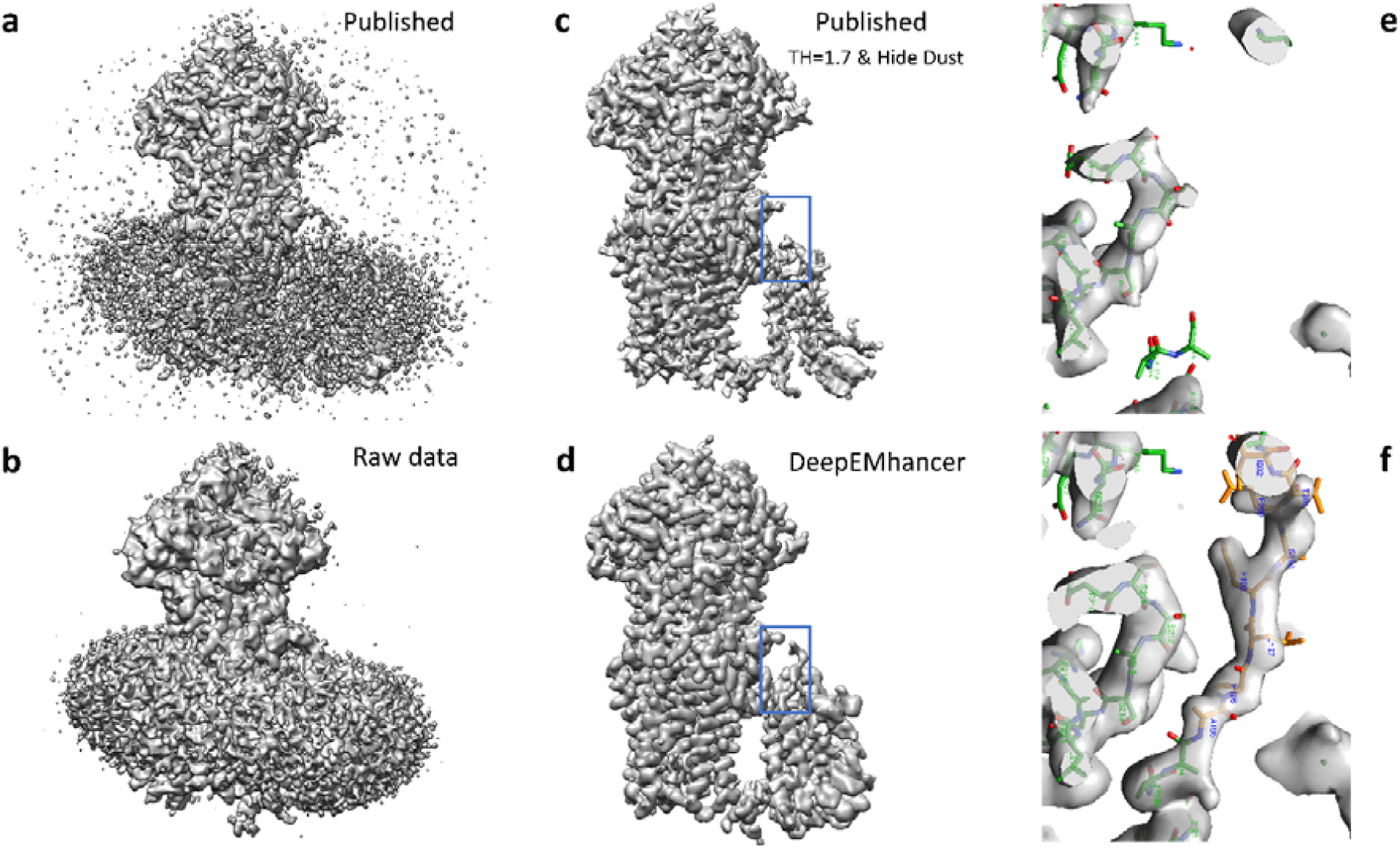
Testing map EMD-7099. **a**, Lateral view of the published map (B-factor sharpened, shown at recommended threshold). **b**, Lateral view of the raw data map obtained from the half maps that was used as input for DeepEMhancer. **c**, Lateral view of the published map after rising the threshold and removing the small connected components so that the signal coming from the lipids was suppressed. As a collateral consequence, some densities corresponding to the protein were also lost. **d**, Lateral view of the map obtained with DeepEMhancer. **e**, Zoom-in of the region marked with a blue box in c. **f**, Zoom-in of the region marked with a blue box in d, in which DeepEMhancer post-processed map, contrary to the published map, shows the densities corresponding to a missing loop in PDB 6bhu chain A. As a result, the residues A195 to I203 have been *de novo* modeled (new residues depicted in yellow, published in green).

#### EMD-4997

The EMD-4997 (Walter et al., 2019) is a medium-high resolution volume (4.0 Å) for a murine epithelial anion transporter. As in the previous example, the overall quality of the map is quite good, yet it presents lower quality regions. Figure 5a shows an overview of the published map, displayed at the recommended threshold, and the map obtained with DeepEMhancer. Although it is true that both the original map and the post-processed map look very similar, it is also true that there exist important differences. Firstly, the map processed with DeepEMhancer is cleaner than the original one. Serve as an example the removal of the artifacts that the published map presents near the elbow of the complex (see Figure 5a, red box). More importantly, there can also be found many regions for which the DeepEMhancer post-processed volume resolves better the different residues of the regions. One of such examples can be found near the N-terminal end of the protein complex. Thus, as it is shown in Figure 5b, the densities that correspond to the strands of the β-sheet are better separated than in the published volume. It is important to notice that this better separation is not a consequence of the employed thresholds, as it is proven by the fact that rising the threshold makes the densities corresponding to the backbone discontinuous before the densities for the two strands separate (see Figure 5b). As a result, we can affirm that the quality of this region has been improved by the usage of DeepEMhancer.

**Figure 5.**
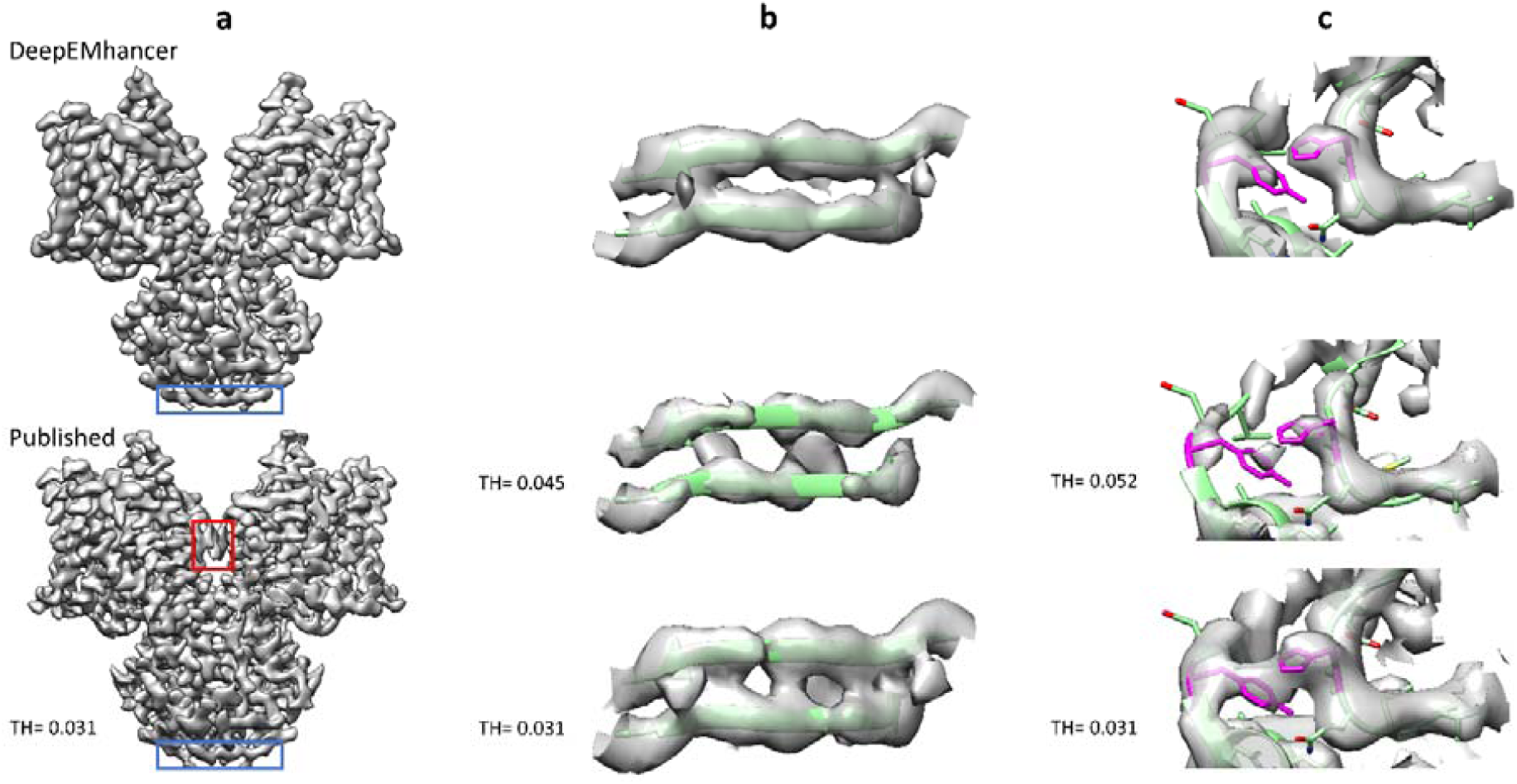
Testing map EMD-4997. **a**, Overview of the published map (B-factor sharpened, shown at the recommended threshold of 0.031), bottom, and the map obtained with DeepEMhancer, top. Red box highlights an artifact that has been automatically removed by DeepEMhancer. Blue box delimits the region showed in b. **b**, Zoom-in of the region marked with a blue box that contains the β-sheet R7-A10, chains A and B. The published volume is shown at the recommended threshold and at the threshold at which the backbone begins to look discontinuous. As it can be appreciated, DeepEMhancer solution resolves better than the published map the two strands of the sheet. **c**, Zoom-in of the region centered at chain B residues H121 and Y361 (colored in magenta). The published volume is shown at the recommended threshold and at the smaller threshold at which the density that connects the two residues disappears. As it can be appreciated, DeepEMhancer post-processed map resolves better than the published map the two residues.

Another similar example is displayed in Figure 5c. In this case, two non-contiguous aromatic residues, Y361 and H121, seem connected in the original map. However, when DeepEMhancer is applied, the densities corresponding to the two residues look separated while the backbone remains continuous.

### Use case EMD-30178 from SARS-CoV-2 RNA-dependent RNA polymerase

In order to further explore the benefits of the DeepEMhancer algorithm we analyzed more deeply the post-processing of EMD-30178 map from Gao et al. (Gao et al., 2020), corresponding to the SARS-CoV-2 RNA-dependent RNA polymerase. The original map presents detailed structure up to 2.9 Å resolution, however, as is often the case in cryo-EM, the resolution of the map is highly heterogeneous. We have chosen this map not only for the importance of this structure in current days but also because of the fact that the heterogeneous quality of the map density presents an ideal case for DeepEMhacer software. As it is shown in Figure 6 a, the application of the algorithm reduces the noise and improves the consistency and depiction of the map. To better illustrate these differences, we have chosen two different regions in chains A and D where the differences between the original and the DeepEMhancer map can be appreciated (Figure 6 b and c). While the density in the original map looks noisy or discontinuous depending on the displayed threshold (Fig. 6 b and c, left and middle panel), the application of the DeepEMhacer software results in a well-defined continuous density where the side chains are nicely depicted (Figure 6 b and c, right panel). This improvement in the map density allowed us to close the loop between residues in the β-sheet V115 to I132 from chain D tracing 3 new residues that were not traced in the original structure (Figure 6 b). The improvement of the density is not only applicable to the edges of the map but it can be also appreciated in its core. Residues H362 to L366 in chain A, traced on the original map where positioned more accurately on the density after map post-processing (Fig 6 c).

**Figure 6.**
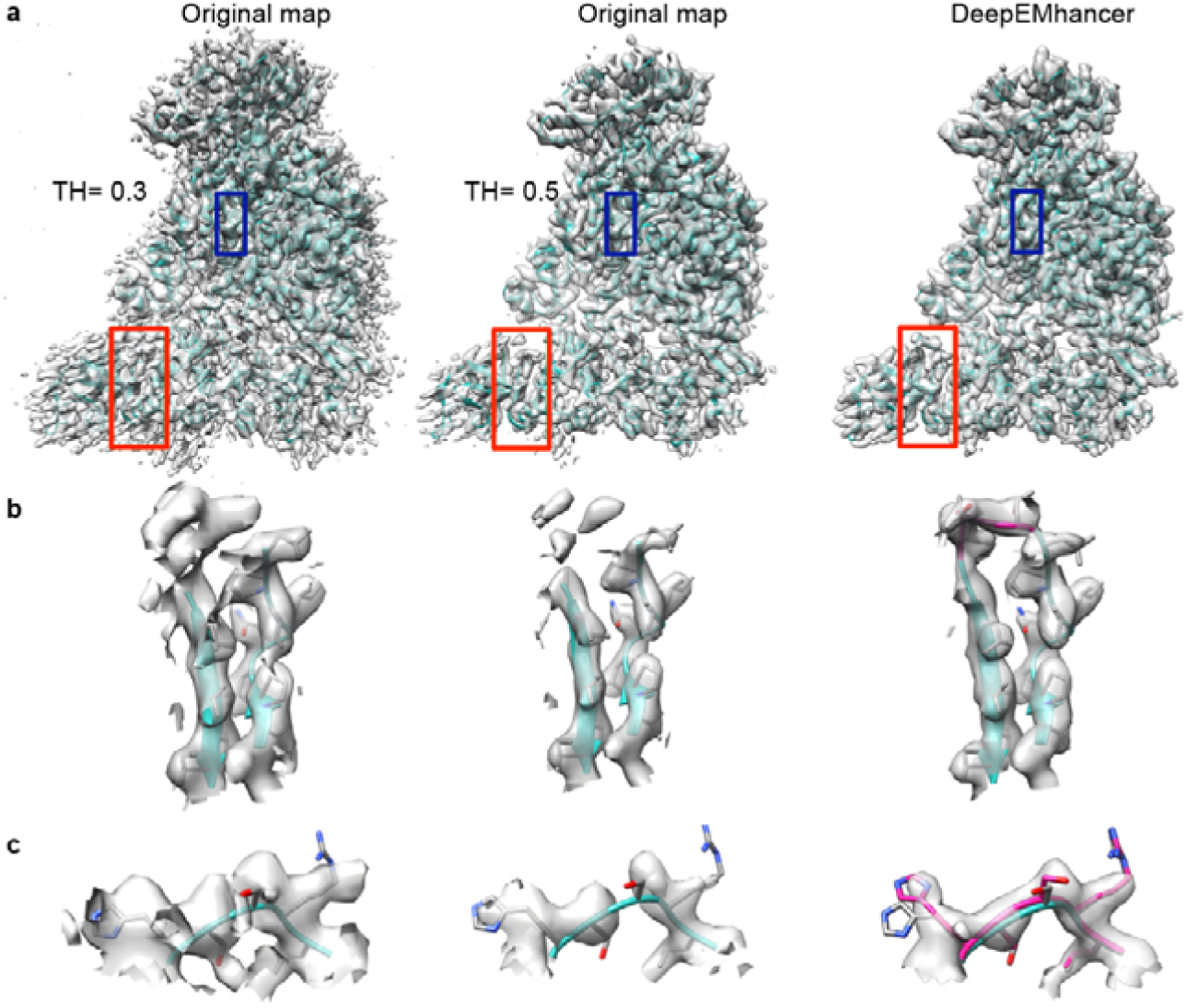
Use case EMD-30178 from SARS-CoV-2 RNA-dependent RNA polymerase. **a**, Overview of the original map displayed with two different thresholds 0.3 (recommended, left) and 0.5 (middle panel) and processed with DeepEMhacer software (right). PDB 7btf is shown in ribbon, red squares designated the zoomed areas in b panel and blue squares the zoomed areas in c. **b**, Zoom-in and extraction of the density from the 3D reconstruction of the original map at different thresholds and DeepEMhacer map corresponding to the red squares in a, chain D from residues V115 to I132. Newly traced residues in the DeepEMhancer map are shown in pink **c**, Zoom-in and extraction of the density from the 3D reconstruction of the original map at different thresholds and DeepEMhacer map corresponding to the blue boxes.

## Discussion

The number of deposited high resolution cryo-EM maps have soared since the beginning of the ‘resolution revolution’. As a result, there is an increasing number of atomic models that are being built using cryo-EM as the primary source of information. However, building atomic models directly from the raw maps is generally not possible. Instead, maps are post-processed in order to enhance the contrast of their high-resolution features.

In this work we have presented Deep cryo-EM Map Enhancer (DeepEMhancer), a new map post-processing method based on deep learning. Trained on pairs of experimental cryo-EM maps and post-processed maps constructed with LocScale using atomic models, DeepEMhancer has learned how to perform a high-quality post-processing operation that reproduces the effects of masking and local sharpening in an automatic fashion.

The performance of our new algorithm has been assessed using a testing set of 20 experimental maps that were not used for training nor during the trial and error process required for its implementation. In all cases, the similarity between the maps obtained from the atomic models and the experimental maps improved after the application of DeepEMhancer. Additionally, we evaluated in detail the performance of DeepEMhancer on two of those maps, showing that, not only DeepEMhancer facilitates the visualization of cryo-EM maps, but also that DeepEMhancer can unveil some details that are not easily recognizable in the raw maps. Finally, we have employed DeepEMhancer on a map of the RNA polymerase of the SARS-CoV 2 virus, improving its quality of the map and the quality of the associated atomic model.

## Methods

### Raw data collection

DeepEMhancer has been trained and evaluated using as input a subset of cryo-EM maps obtained from the EMDB (Lawson et al., 2015) that meet the following requirements: 1) resolution better than 7 Å; 2) have one and only one atomic model associated; 3) correlation between the atomic model and the map better than 0.6 and 4) half maps available. As a result, an original list of 415 maps was compiled. However, this initial list is highly redundant and, in order to avoid biases in both the training and evaluation procedures, this list was further filtered to reduce its redundancy, (see subsection *Redundancy control*). Finally, after a visual inspection aimed at removing problematic cases that survived to the automatic filtering procedure, a total amount of 151 maps, with an average reported resolution of 3.8 Å, was selected.

Since the main objective of DeepEMhancer is to perform a sharpening-like post-processing transformation, it is important to ensure that the maps used in this study were not previously sharpened. Given the fact that most of the maps deposited in EMDB are sharpened and many are also masked, we decided to employ only the half-maps available in EMDB (condition number 4). Due to the lack of an appropriate searching tool in EMDB and a file name convention, we had to analyze all the map file names included in the database looking for the substring “half” to recover the half maps. Full maps were obtained averaging respective half maps.

As learning targets, we employed the output generated by LocScale using as input the aforementioned maps and their associated atomic models. Additionally, the output maps were tightly masked using as masks the maps simulated from the atomic models after a thresholding operation. Although it is true that the simulated maps could be directly employed as targets, we discarded this alternative for two reasons. The first one is empirical: we obtained better results when targets were produced with LocScale, probably because the input and target maps, although different, they still share some similar properties such as intensity ranges or local quality, which are not necessarily preserved when using simulated maps as targets. The other reason is that we wanted to reproduce the state-of-the-art local sharpening effect and not a new type of post-processing that could not be compatible with downstream atomic modelling tools.

### Data preparation

Due to the fact that the monomers (amino acids, nucleotides…) that compose the macromolecules have fixed size but the deposited maps vary in voxel size, both the input and the target maps were resampled to 1 Å/voxel size with the aim of facilitating the learning process. After that, the intensity of each volume was normalized using the classical cryo-EM approach by which the map noise statistics are forced to adopt a fixed mean and standard deviation (0 and 0.1 respectively). Finally, due to GPU size limitations, the maps were chunked into 64×64×64 cubes, the maximum size that our computing systems were able to efficiently manage. As a result, more than 70k volume cubes, including both signal cubes and noise-only cubes were used for training.

### Redundancy control

In order to perform the train/test/validation split used to develop and evaluate our method, it is important to consider that the universe of proteins is highly redundant and that the EMDB entries are even more redundant. Serve as an example the case of the ribosome, that supposes ∼10% of the all EMDB entries. Thus, in order to avoid an over-optimistic performance estimation, we have ensured that the train, test and validation sets are mutually exclusive in the sense that their intersections are empty under a certain equivalence criterion. Particularly, we consider that two EMDB entries are equivalent if they share one sequence that belongs to the same 30% sequence identity cluster. Similarly, with the aim of eliminating potential bias in the evaluation, we have guaranteed that only one member per cluster is included in testing and validation sets. On the contrary, we have relaxed our quite strict redundancy control policy in the training set allowing up to five cluster representatives in an attempt to increase the size of this set. This decision is founded on the fact that even maps of the same exact protein may present different statistics due to the intrinsic variability of cryo-EM reconstruction workflows and thus, limiting their presence in the training set may difficult the generalization of the neural network. As a result, a list of 110, 21 and 20 maps were used for training, validation and testing respectively. The full list of the EMDB entries used can be found in Supplementary Material.

### Neural network architecture

We have employed a 3D U-net-like neural network (Ronneberger et al., 2015) as a regression model for the estimation of post-processed maps. Our neural network consists of three downsampling blocks and three upsampling blocks with skip connections. Each block contains three convolutional layers followed by group normalization (Wu and He, 2020) and PRelu activation (He et al., 2015). The number of filters for each block is 3×32, 3×64 and 3×128 respectively. Downsampling is carried out using strided convolutions and upsampling is performed via transposed convolution. See Supplementary Material for additional details.

### Neural network training

Our neural network was trained using stochastic gradient descent with a batch size of 8 cubes. Initial learning rate was set to 10^−3^ and decreased by a factor of 0.5 when the validation loss did not improve during 5 epochs. As loss function, mean absolute error was employed. Data augmentation, consisting in random 90° rotations, gaussian blurring and patch corruption was applied to the training data.

### Neural network inference

In order to perform volume post-processing, the input volume is pre-processed as described in the *Data preparation* subsection. Then, the resized and normalized volume is chunked into overlapping cubes of size 64×64×64 with strides of 16 voxels. Each cube is individually processed by the trained neural network, yielding post-processed cubes. After that, the post-processed cubes are re-assembled into the final volume averaging the overlapping parts. Finally, the processed volume is resized to the size of the original volume, thus, showing the correct sampling rate value.

### Evaluation

With the aim of guiding the cross-validation process, we computed the correlation coefficient between the maps produced by DeepEMhancer and the maps used as learning targets (masked LocScale post-processed maps). Once the final model was selected, the quality of DeepEMhancer predictions were assessed comparing the input and processed maps against the reference maps obtained from the atomic models. Specifically, we computed the Fourier Shell Correlation coefficient (FSC) between them and we estimated the resolution using 0.5 as threshold. Due to the fact that DeepEMhancer performs a non-conventional post-processing operation, including masking and enhancement operations, in order to disentangle the two effects, the FSC was also computed after masking the maps to compare with a tight mask derived from the atomic model.

As a complementary metric, we also applied DeepRes (Ramírez-Aportela et al., 2019) over the input and processed maps. DeepRes is a deep learning-based local resolution method that, contrary to others, is sensitive to the sharpening process and thus, it can provide an alternative estimation of the post-processing effect.

Finally, for comparison purposes, we repeated the FSC and DeepRes experiments using the Relion postprocessing program (Kimanius et al., 2016; Zivanov et al., 2018). As Relion automatic masking is very simple, in order to make the comparison more interesting, we decided to execute the postprocessing algorithm using the mask derived from the atomic models. Similarly, since the automatic determination of the B-factor can produce worse results than a manually selected one, in addition to the maps computed using an automatically determined B-factor by Relion, we also considered the sharpened map deposited in EMDB.

### EMD-30178 map evaluation and atomic model modification

DeepEMhancer was applied to the half maps deposited in EMDB entry EMD-30178. The original and post-processed maps were visually inspected using Coot (Emsley and Cowtan, 2004) and chimera (Pettersen et al., 2004), and chosen regions on the 7btf PDB were newly built or modified using Coot.

## Data availability

DeepEMhancer is available at https://github.com/rsanchezgarc/deepEMhancer and as an Xmipp protocol for Scipion v3. Post-processed map examples are available at http://campins.cnb.csic.es/deepEMhancer/examples.

## Supporting information

Supplementary sections

## Acknowledgments

This work is supported by the the Comunidad de Madrid through grant CAM (S2017/BMD-3817), the Spanish Ministry of Economy and Competitiveness (BIO2016-76400-R) and the Spanish Ministry of Science and Innovation through the call 2019 Proyectos de I+D+i - RTI Tipo A (PID2019-108850RA-I00). J.V. acknowledges economical support from the Ramón y Cajal 2018 program (RYC2018-024087-I). R.S.G. is recipient of an FPU fellowship.

## Author Contributions

Conceptualization, J.V., C.O.S.S., R.S.G.; Methodology, R.S.G., J.V. and C.O.S.S.; Software implementation, R.S.G., J.G.; Evaluation, R.S.G., J.G., A.C.; Writing, R.S.G., A.C., J.V., C.O.S.S. and J.M.C.; Supervision, J.V., C.O.S.S.; Funding Acquisition, J.M.C, J.V.

## Notes

### Competing Interest Statement

The authors have declared no competing interest.

### Summary of Updates

Figures updated. Shorter summary

