## Supplementary sections for "DeepEMhancer: a deep learning solution for cryo-EM volume post-processing"

### Neural network architecture

**Table ST1**. Neural network architecture

| Layer | Type | Parents | #Kernels | Stride | Kernel size | Output shape |
|---|---|---|---|---|---|---|
| 1 | Conv3d+GN+PReLU | Input | 32 | 1 | 5 | 64x64x64x32 |
| 2 | Conv3d+GN+PReLU | 1 | 32 | 1 | 5 | 64x64x64x32 |
| 3 | Conv3d+GN+PReLU | 2 | 32 | 2 | 5 | 32x32x32x32 |
| 4 | Conv3d+GN+PReLU | 3 | 64 | 1 | 5 | 32x32x32x64 |
| 5 | Conv3d+GN+PReLU | 4 | 64 | 1 | 5 | 32x32x32x64 |
| 6 | Conv3d+GN+PReLU | 5 | 64 | 2 | 5 | 16x16x16x64 |
| 7 | Conv3d+GN+PReLU | 6 | 128 | 1 | 5 | 16x16x16x128 |
| 8 | Conv3d+GN+PReLU | 7 | 128 | 1 | 5 | 16x16x16x128 |
| 9 | Conv3d+GN+PReLU | 8 | 128 | 2 | 5 | 8x8x8x128 |
| 10 | Conv3d+GN+PReLU | 9 | 128 | 1 | 5 | 8x8x8x128 |
| 11 | Concat +Conv3d_trans | 10 | 128 | 2 | 5 | 16x16x16x128 |
| 12 | Conv3d+GN+PReLU | 10 & 7 | 128 | 1 | 5 | 16x16x16x128 |
| 13 | Conv3d+GN+PReLU | 12 | 128 | 1 | 5 | 16x16x16x128 |
| 14 | Conv3d+GN+PReLU | 13 | 128 | 1 | 5 | 16x16x16x128 |
| 15 | Concat +Conv3d_trans | 14 & 4 | 64 | 2 | 5 | 32x32x32x64 |

| | | | | | | |
|---|---|---|---|---|---|---|
| 16 | Conv3d+GN+PReLU | 15 | 64 | 1 | 5 | 32x32x32x64 |
| 17 | Conv3d+GN+PReLU | 16 | 64 | 1 | 5 | 32x32x32x64 |
| 18 | Conv3d+GN+PReLU | 17 | 64 | 1 | 5 | 32x32x32x64 |
| 19 | Concat +Conv3d_trans | 18 & 1 | 32 | 2 | 5 | 64x64x64x32 |
| 20 | Conv3d+GN+PReLU | 19 | 32 | 1 | 5 | 64x64x64x32 |
| 21 | Conv3d+GN+PReLU | 20 | 32 | 1 | 5 | 64x64x64x32 |
| 22 | Conv3d+GN+PReLU | 21 | 32 | 1 | 5 | 64x64x64x32 |
| 23 | Conv3d_trans | 22 | 16 | 2 | 5 | 128x128x128x16 |
| 24 | Conv3d+GN+PReLU | 23 | 8 | 2 | 5 | 64x64x64x8 |
| 25 | Conv3d | 24 | 1 | 1 | 5 | 64x64x64x1 |

Total number of parameters: 51,119,889

#### Performance on the testing set

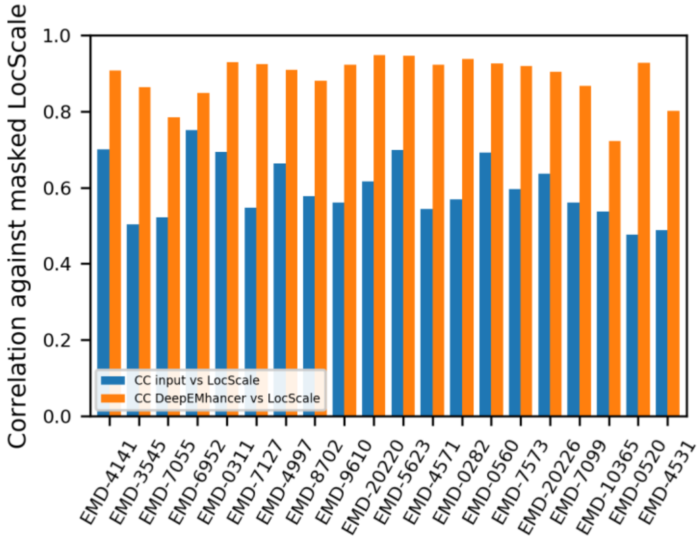

**Figure S1**. Correlation coefficient for the input maps of the testing set before and after the treatment with DeepEMhancer

#### Visual inspection of testing maps

*EMD-7055*

The EMD-7055 (Tenthorey et al., 2017) is a medium resolution volume of the NAIP5-NLRC4-flagellin inflammasome. For the purposes of this article, the main interesting aspect of this volume is the apparent poor performance of DeepEMhancer according to main text Figure 1. One of the reasons behind this behaviour is the fact that a mask was applied to only three subunits at the last stages of the refinement process. As a result, the volume contains signal for both masked and unmasked subunits although their intensity levels vary severely. Consequently, our neural network has tried to restore both the originally masked and unmasked regions, and thus, the results are not as good as in the other cases. However, when the volume is carefully pre-processed in order to remove those unmasked subunits while preserving the normalization constraints, non-negligible improvements were observed. Secondly, another important reason for the poor measured metrics is the fact that the atomic model (PDB 6b5b) was obtained by means of rigid body fitting of an homology model instead of being traced, thus, the agreement between the atomic model and the density map is far from being perfect. As a result, the resolution estimates computed using the atomic model as reference are not too accurate.

Figure S2 shows the overall aspect of the post-processed volume compared to the raw and the B-factor-sharpened ones. As can be appreciated in Figure S2 A, the map produced by DeepEMhancer is much cleaner than the B-factor processed map. More importantly, although the level of detail in the core of the protein is similar, in the outer part of the protein, the B-factor sharpened map presents broken densities that look continuous in the map obtained with DeepEMhancer, thus facilitating the map interpretation.

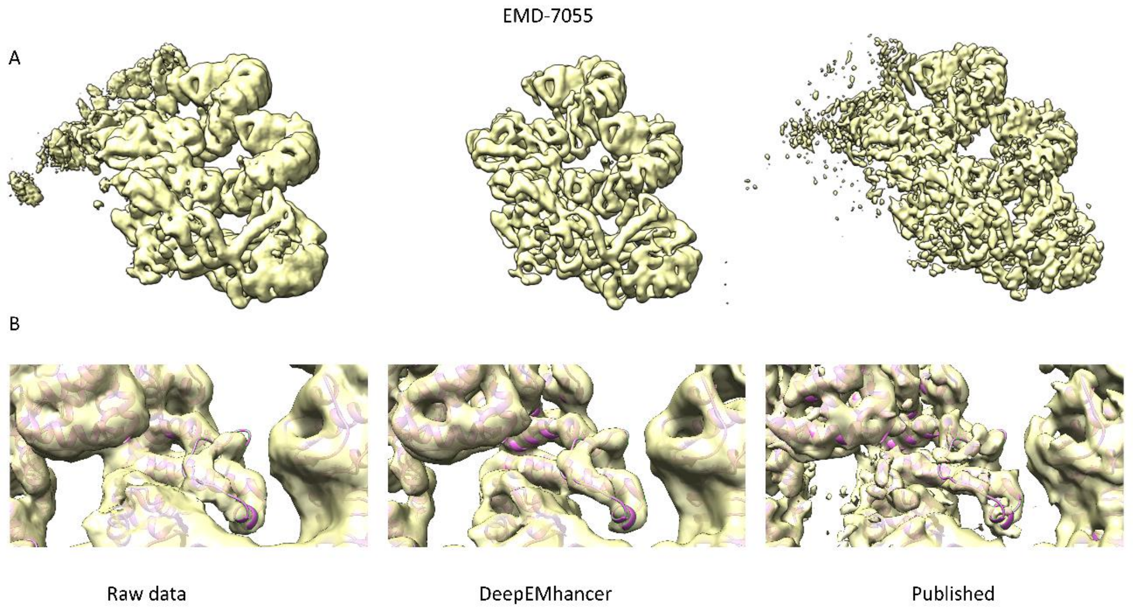

**Figure S2**. DeepEMhancer post-processed volume for EMD-7055. A, overview of the raw data map, the post-processed map and the B-factor corrected map. B, zoom-in of a region containing the loop Q514-S533, which looks cleaner and better resolved in the DeepEMhancer post-processed map compared to the raw data and sharpened maps.

### List of EMDB entries used in this work

#### Training

EMD-0026
EMD-0038
EMD-0071
EMD-0093
EMD-0094
EMD-0132
EMD-0234
EMD-0244
EMD-0408
EMD-0415
EMD-4288
EMD-0452
EMD-0490
EMD-0492
EMD-0500
EMD-0501
EMD-0552
EMD-0567
EMD-0589
EMD-0592
EMD-0665
EMD-0776
EMD-10049

EMD-10069
EMD-10100
EMD-10105
EMD-10106
EMD-10134
EMD-10273
EMD-10279
EMD-10324
EMD-10333
EMD-10418
EMD-10534
EMD-10585
EMD-10595
EMD-10617
EMD-20145
EMD-20146
EMD-20189
EMD-20234
EMD-20249
EMD-20254
EMD-20259
EMD-20270
EMD-20271
EMD-20352
EMD-20521
EMD-20986
EMD-21012
EMD-21107
EMD-21144
EMD-21391
EMD-3661
EMD-3662
EMD-3802
EMD-3885
EMD-3908
EMD-4032
EMD-4073
EMD-4074
EMD-4079
EMD-4148
EMD-4162
EMD-4192
EMD-4214
EMD-4241
EMD-4272
EMD-4401
EMD-4404
EMD-4429
EMD-4588
EMD-4589
EMD-4593
EMD-4728
EMD-4746
EMD-4748
EMD-4759

EMD-4888
EMD-4889
EMD-4890
EMD-4907
EMD-4917
EMD-4918
EMD-4941
EMD-4983
EMD-6479
EMD-7009
EMD-7041
EMD-7065
EMD-7090
EMD-7334
EMD-7335
EMD-7770
EMD-7869
EMD-8437
EMD-8438
EMD-8911
EMD-8958
EMD-8960
EMD-9111
EMD-9258
EMD-9259
EMD-9891
EMD-9931
EMD-9934
EMD-9935
EMD-9939
EMD-9941
EMD-9695

Validation

EMD-0193
EMD-0257
EMD-0264
EMD-0499
EMD-10401
EMD-20133
EMD-20449
EMD-20508
EMD-20849
EMD-4611
EMD-4646
EMD-4733
EMD-4789
EMD-6847
EMD-7133
EMD-7882
EMD-8069

EMD-9112
EMD-9298
EMD-9374
EMD-9664

Test

EMD-0282
EMD-0311
EMD-0520
EMD-0560
EMD-10365
EMD-20220
EMD-20226
EMD-3545
EMD-4141
EMD-4531
EMD-4571
EMD-4997
EMD-5623
EMD-6952
EMD-7055
EMD-7099
EMD-7127
EMD-7573
EMD-8702
EMD-9610
